# A single-cell atlas of the murine pancreatic ductal tree identifies novel cell populations with potential implications in pancreas regeneration and exocrine pathogenesis

**DOI:** 10.1101/2024.02.26.582044

**Authors:** Ángel Fernández, Joan Casamitjana, Adrián Holguín-Horcajo, Katarina Coolens, Loris Mularoni, Lorenzo Pasquali, Jennifer M. Bailey-Lundberg, Ilse Rooman, Yue J. Wang, Meritxell Rovira

## Abstract

**Background and aims:** Pancreatic ducts form an intricate network of tubules that secrete bicarbonate and drive acinar secretions into the duodenum. This network is formed by centroacinar cells, terminal, intercalated, intracalated ducts, and the main pancreatic duct. Ductal heterogeneity at the single-cell level has been poorly characterized; therefore, our understanding of the role of ductal cells in pancreas regeneration and exocrine pathogenesis has been hampered by the limited knowledge and unexplained diversity within the ductal network.

**Methods:** We used scRNA-seq to comprehensively characterize mouse ductal heterogeneity at single-cell resolution of the entire ductal epithelium from centroacinar cells to the main duct. Moreover, we used organoid cultures, injury models and pancreatic tumor samples to interrogate the role of novel ductal populations in pancreas regeneration and exocrine pathogenesis.

**Results:** We have identified the coexistence of 15 ductal populations within the healthy pancreas and characterized their organoid formation capacity and endocrine differentiation potential. Cluster isolation and subsequent culturing let us identify ductal cell populations with high organoid formation capacity and endocrine and exocrine differentiation potential *in vitro*, including Wnt-responsive-population, ciliated-population and FLRT3^+^ cells. Moreover, we have characterized the location of these novel ductal populations in healthy pancreas, chronic pancreatitis, and tumor samples, highlighting a putative role of WNT-responsive, IFN-responsive and EMT-populations in pancreatic exocrine pathogenesis as their expression increases in chronic pancreatitis and PanIN lesions.

**Conclusions:** In light of our discovery of previously unidentified ductal populations, we unmask the potential roles of specific ductal populations in pancreas regeneration and exocrine pathogenesis.

## INTRODUCTION

Pancreatic ducts form an intricate network of tubules, the first cells of the ductal epithelium touching the acini are the centroacinar cells (CACs), located at the tip of terminal/intercalated ducts that will later join to form larger intralobular ducts that fuse into interlobular ducts to empty acinar secretions into the main duct. The main duct merges with the common bile duct and opens into the duodenum, through the duct of Wirsung. In addition, there is an accessory duct, named duct of Santorini, that joins the main duct in the head of the pancreas^1^. Ductal cells secrete an alkaline mucus fluid rich in bicarbonate to neutralize the acidic chyme of the stomach. Moreover, ductal cells play a key role in the development of pancreatic exocrine pathologies, such as cystic fibrosis, acute and chronic pancreatitis, and pancreatic cancer (PDAC)^1^. Notably, well-differentiated PDAC carries the morphological appearance of ductal structures expressing ductal markers^2^. Thus, by histology alone, the putative cell of origin was long thought to be ductal^3^. However, recent studies demonstrated that both acinar and ductal cells can be transformed upon *KRAS* mutations^4^.

Tissue-resident progenitors have been studied in highly proliferative and regenerative tissues, such as the intestine^5^, where progenitors comprise 5-8% of the tissue^6^. However, the pancreas has low proliferation rates and limited capacity for regeneration, thus the reservoir of progenitors, if any, is likely modest. Ductal cells, in the mammalian pancreas, have attracted most of the attention as potential sources of new β cells for several reasons. 1) Several studies observed endocrine cells near to or embedded in the ducts during growth or regeneration^7,8^; 2) Ductal cells and endocrine cells share a common progenitor during development. Neurogenin3^+^ cells within these progenitors delaminate and differentiate into endocrine cells while the remaining duct-like complexes differentiate into mature ducts^9,10^. 3) Moreover, lineage tracing of ductal cells in zebrafish demonstrated that they are progenitors in adult tissue^11^. Therefore, it stands to reason that ductal cells may play a role in pancreas regeneration, although it is a highly controversial topic as several lineage tracing studies using pan-ductal markers showed no β-cell neogenesis from ducts (reviewed^12,13^). 4) Finally, ductal cells acquire cellular plasticity *in vitro* as they are uniquely able to form organoids^14^; an ability solely displayed by adult progenitors^15^.

Although the ductal network is a complex system, it is mostly seen as a homogeneous population, and little is known about ductal heterogeneity in mammals and the implications of different ductal populations in exocrine pathogenesis and regeneration. Single-cell technologies now allow the investigation of the transcriptome of individual cells, dissecting tissue, and cell heterogeneity beyond what was possible with bulk approaches. Importantly, ductal pancreatic cells have been vastly underrepresented in single-cell analysis for several reasons: i) most of the studies have centered their interest in isolated islets, containing few ducts^16^; ii) digestive enzymes cause degradation of cells and RNA; iii) ducts represent ∼10%^1^ of the gland, but they are refractory to disaggregation; thus, low heterogeneity has been described^17,18^; iv) snRNA-seq of frozen human pancreas has circumvented digestive enzyme activity^19^ but, again, ducts are underrepresented.

Here, we investigated ductal heterogeneity of the entire ductal network using a mouse transgenic line where GFP expression is driven by *Sox9* promoter, labeling all ductal cells^20^. We observed increased capacity of organoid formation ability in medium-big duct-derived organoids with lower Sox9 expression when compared to small ducts. Moreover, the capacity to give rise to endocrine progenitors, insulin-producing and somatostatin-producing cells was significantly higher in medium-big duct-derived organoids. Furthermore, we have comprehensively characterized mouse ductal heterogeneity at single-cell resolution, highlighting the coexistence of 15 ductal clusters. Importantly, we have also identified surface markers allowing the isolation of several novel ductal populations and assess their plasticity in functional studies in organoid cultures showing that ductal populations located in medium-big size ducts have higher progenitor capacity defined by organoid formation efficiency and increased endocrine/exocrine differentiation. Finally, we have characterized tissue expression and localization of these previously unidentified ductal populations in mouse adult pancreas, chronic pancreatitis, and pancreatic tumors.

## MATERIAL AND METHODS

### Mice

*Tg(Sox9-EGFP)EB209Gsat* was obtained from J. Ferrer laboratory (CRG). Experimental procedures were approved by the Animal Experimentation Ethics Committee at IDIBELL (approval #AR18009). 8-16-week-old mice were used in this study. Mice were housed at IDIBELL animal facility under SPF conditions in accordance with the institutional guidelines and ethical regulations. Detailed genotyping primers in **Supplementary_Table 5**.

### Chronic pancreatitis

Chronic pancreatitis mice were housed under standardized SPF conditions and experiments were approved by the Ethical Committee for Animal Experiments of the Vrije Universiteit Brussel and carried out according to the national guidelines on animal experimentation. For the induction of chronic pancreatitis, 8-12-week-old C57BL/6 mice were treated with 125µg/kg caerulein following the protocol described by Grimont et al.^21^. Further details in supplementary methods.

### PDAC models

Two mouse models of PDAC have been used in our studies. Firstly, Pdx-1-Cre mice were crossed with LSL-Trp53^R172H/+^ and LSL-Kras^G12D/+^ mice to obtain LSL-Kras^G12D/+^;LSL-Trp53^R172H/+^;Pdx-1-Cre animals, in accordance with institutional ethical guidelines and approved by St. Vincent’s Hospital Animal Ethics Committee, Sydney, Australia (approval #16/02). The second model used was specific to trace ductal-derived tumors, as previously published^22^, *LSL-Kras^G12V^;* Trp53^R172H/+^; Hnf1bCreERT2 mice were treated with 5mg of tamoxifen to induce PDAC.

### Duct isolation

Mouse pancreas digestion followed previously published protocols^23^ with minor modifications. Finally whole ducts were hand-picked and sorted by size under a fluorescent stereoscope. Further details in supplementary methods.

### Single-cell prep

Small, medium, and big ducts were further digested with tryPLE Express for 5-10min (depending on the size of the ducts) at 37°C, vigorously shaking the digestion mix every 2-3 min. Purified single ductal cells were resuspended in FACS buffer. Further details in supplementary methods.

### FACS analysis

For FACS analysis, antibody incubation was performed following manufacturer’s instructions (**Supplementary_Table 6 and Supplementary_methods**).

FACS was performed using a Beckman Coulter High-Speed Cell Sorter Moflo-XDP. Flow derived data were analysed using the Kaluza 2.1 software.

### Organoid culture

FACS-sorted ductal cells (GFP+) or whole ducts were embedded in Matrigel at a density of 1,000-5,000 cells in 24-well plates and cultured in pancreatic organoid expansion medium **(Supplementary_Table 6 and Supplementary_methods).**

### Endocrine differentiation

Two days after organoid passage, organoids were induced to differentiate with endocrine differentiation medium **(Supplementary_Table 6 and Supplementary_methods)**

### Immunostainings

Staining details are included in supplementary methods. Antibodies are shown in **Supplementary_Table 6**. Confocal images were captured by LeicaTCS_SP5 confocal. Epifluorescent images were acquired on a ZeissAxio_ObserverZ1_Apotome inverted fluorescent microscope. Organoid brightfield images were taken on a LeicaZ16 APO stereomicroscope. Organoid sizes were measured using ImageJ/FIJI. Quantification of marker expression was performed using the Imaris9.6 software.

### RNA isolation and qPCR

RNA isolation of primary cells or organoids was performed following manufacturer’s instructions using RNeasy Mini kit (Qiagen). Extracted RNA was DNAse treated, and reverse transcribed using the Roche Transcriptor First Strand cDNA Synthesis kit. qPCR samples were prepared with SYBR Green and marker specific primers. qPCRs were ran using Applied Biosystems 7900HT Real-Time PCR machine. Primers listed in **Supplementary_Table 5**.

### Sample preparation for scRNA-seq

Ten thousand Sox9:eGFP^+^ cells from each ductal fraction were sorted from a pull of 3 male mice (8-12 weeks) in two independent experiments. Library Preparation of 3’ mRNA was prepared following 10xGenomics manufacturing instructions. Sequencing was carried out as paired-end (PE50) using NovaSeq6000.

### scRNA-seq analysis

Data quality control, filter, dimension reduction, clustering, and differential expression analysis were performed with Seurat V4.1.1 pipeline previously described^24^. Briefly, cells were filtered with nFeature_RNA>200, nFeature_RNA<6000, percent.mt<5, and nCount_RNA<50000. Sctransform was used to normalize the data and regress out potential sample batch effect. RunPCA, RunUMAP, FindNeighbors, and FindClusters (resolution 0.6) were then sequentially applied to perform dimension reduction and cell clustering. FindAllMarkers were then called with min.pct=0.1 and logfc.threshold=0.25 to identify markers for each clusters.

### Statistical analysis

Unless specified in the figure legend, all the p-values were calculated using GraphPad PRISM10 with the following significance: n.s. p-value>0.05; *p-value<0.05; **p-value<0.01; ***p-value<0.001; ****p-value<0.0001. Statistical tests used for each experiment can be found in figures legends.

## RESULTS

### Different organoid formation potential of ductal compartments fractionated by size

Ductal cells can form organoids in culture upon the addition of developmental cues in the medium mimicking the niche^14^, although it is not known if all pancreatic ductal cells display equal organoid formation capacity. Thus, we investigated if ductal cells from different compartments of the ductal tree, including CACs, terminal ducts (TD), intracalated ducts (IAD), intercalated ducts (IED), and main duct (MD) display different capacities for organoid formation **(Figure 1A-B).** Upon pancreas digestion, we isolated small (S<75 µm, including CACs and TD), medium (M:75-250 µm, including small IAD and small IED), and big ducts (B>250 µm, including big IAD and big IED and MD) guided by a Sox9:eGFP transgenic line **(Figure 1B-C).** This line recapitulates Sox9 expression in all tissues, including the pancreas^20^, where the entire ductal epithelium expresses Sox9^10^ **(Figure 1C-D).** While all the fractions formed organoids, medium-big ducts form bigger organoids in a short period of time **(Figure 1E)**.

**Figure 1:**
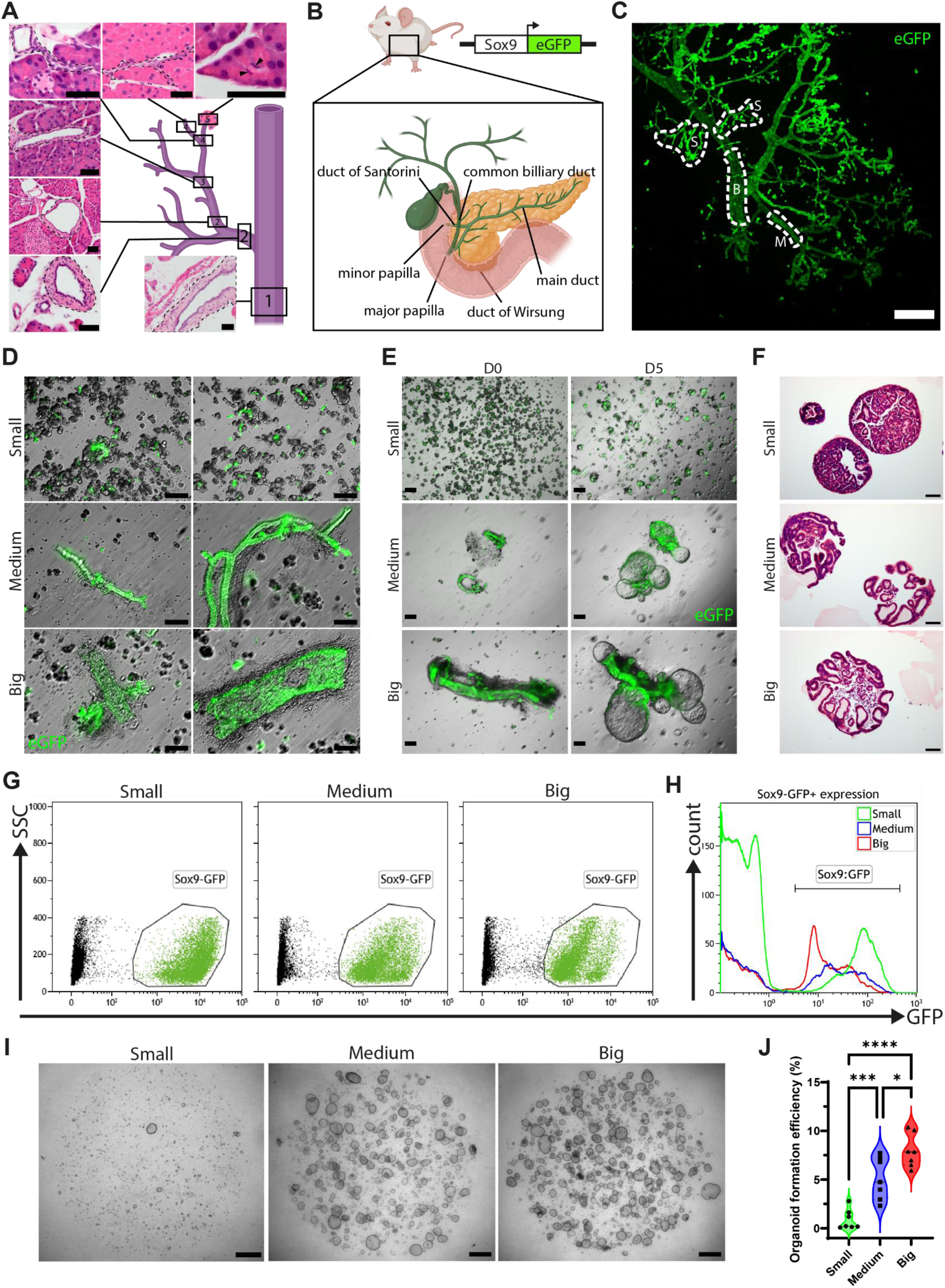
Organoid formation potential of ductal compartments. **A)** Drawing and representative images (hematoxylin-eosin) of the pancreatic ductal tree, showing main duct (1), interlobular (2), intralobular (3), intercalated/terminal (4) ducts and CACs (5). Dashed lines indicate the ducts and arrowheads CACs. Scalebars=50μm. **B)** Drawing of the Sox9:eGFP transgenic mouse. **C)** Maximum projection of lobular units of a Sox9:eGFP pancreas intercalated ducts and CACs (S, small), intralobular (M, medium), and interlobular (B, big) ducts highlighted. Scalebar=200μm. **D)** Representative brightfield images of small, medium, and big ducts hand-picked following digestion. Enriched fractions may have acini (A), blood vessels (BV), and/or stroma. Scalebar=100μm. **E)** Ducts cultured at day 0 and 5 days in organoid expansion medium. Scalebars=200μm. **F)** Representative FACS plots of Sox9:eGFP^+^ cells isolated from small, medium, and big ducts. **G)** Representative FACS histogram showing Sox9:eGFP levels that increase as the size of the ducts diminishes. **I)** Images of organoids plated at same cell density after one week in culture. Scalebars=100μm. **J)** Violin plot showing organoid formation efficiency (1-way ANOVA, Tukey *post hoc*, n=7).

### Bigger ducts display higher organoid formation capacity

To ensure that the different organoid formation ability is not due to i) a higher number of cells in bigger ducts, ii) signaling from vessels, connective tissue, extracellular matrix, or mesenchyme, enriched in big ducts, or iii) counterproductive effect of acinar cells enriched in small duct fractions **(Figure 1D-E),** we isolated Sox9:eGFP^+^ cells by FACS of each fraction **(Figure 1F).** The isolation of ductal cells highlighted Sox9 heterogeneous expression, showing a reduction of Sox9:eGFP intensity as ductal size increases **(Figure 1F-G).** Isolated ductal cells were plated at same density to test organoid formation capacities. Big and medium ductal-derived organoids showed higher organoid formation efficiency in number and size **(Figure 1H-I and Supplementary_Figure 1A)**; 0.89%±1.03 for small, 5.11%±2.16 for medium and 7.93%±1.73 for big ductal-derived organoids.

### Medium-Big ductal-derived organoids display increased differentiation potential into endocrine lineages

Organoids expressed classical ductal markers independent of the duct of origin **(Figure 2A)**. However, we observed significant heterogeneity, while *Hnf1β* expression was similar in all organoids, *Sox9* and *Spp1* expression was lowest in bigger ducts. Genes most highly expressed in big ducts-derived organoids included *Krt19* and *Onecut2* **(Figure 2A)**. Further characterization of ductal markers by immunofluorescence in organoids showed a high degree of heterogeneity **(Supplementary_Figure 2A-B)**. These results suggest the concurrence of diverse differentiation states or the coexistence of different ductal populations in the original preparation.

**Figure 2:**
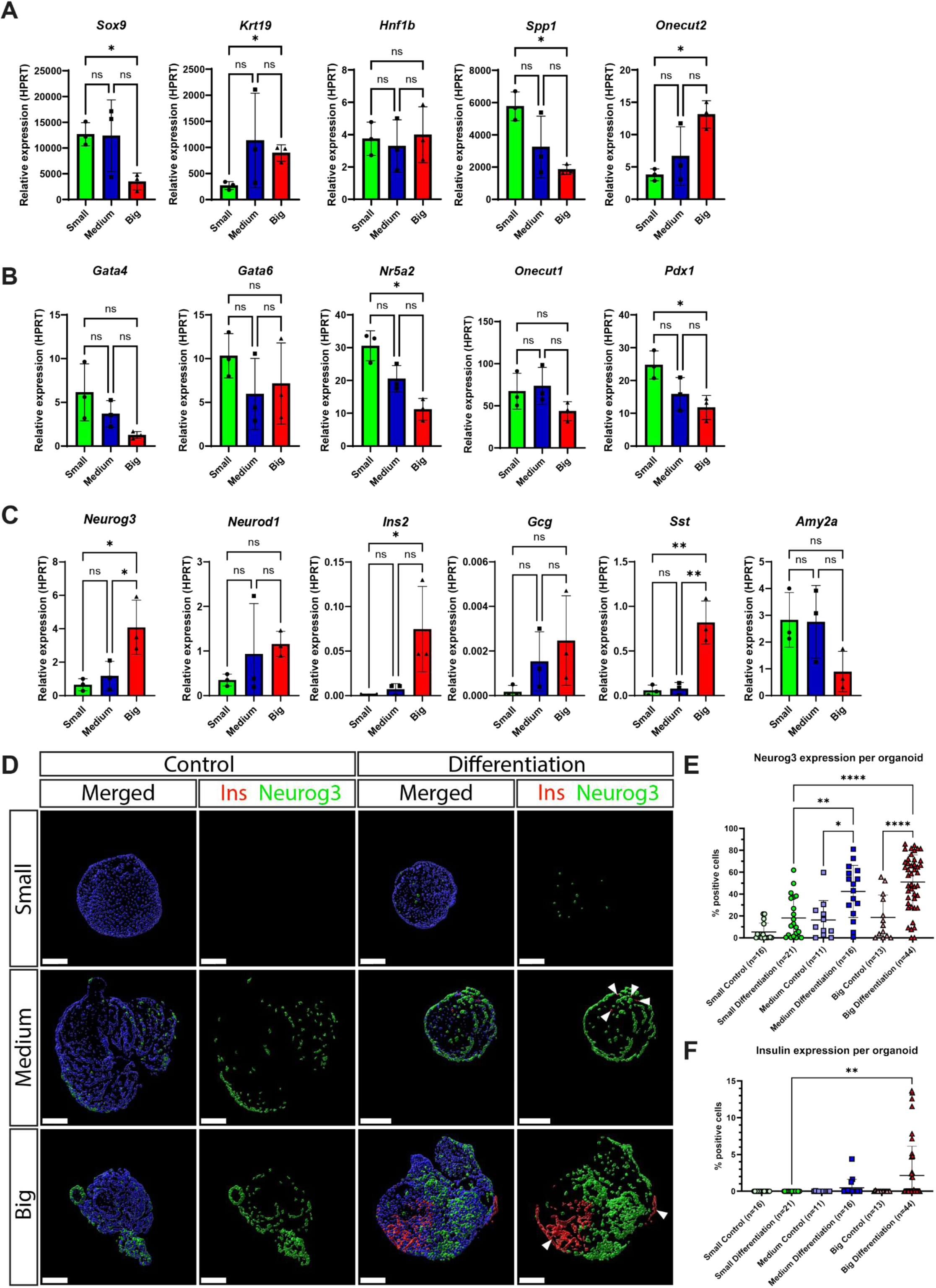
Organoid differentiation potential into endocrine progenitors and insulin-producing cells. **A)** mRNA expression of ductal markers (*Sox9, Krt19, Hnf1b, Spp1 and Onecut2*) by qPCR on small, medium, and big duct-derived organoids (Brown-Forsythe and Welch ANOVA, Dunnett’s T3 *post hoc*, n=3) **B)** mRNA expression analysis of genes expressed in multipotent pancreatic and bipotent progenitors (*Gata4, Gata6, Nr5a2, Onecut1 and Pdx1*) on pancreatic derived organoids (Brown-Forsythe and Welch ANOVA, Dunnett’s T3 *post hoc*, n=3). **C)** Endocrine differentiation was assessed by qPCR analysis of *Neurog3, NeuroD1*, *Ins2, Gcg, Sst and Amy2a* mRNA levels on duct-derived organoids upon differentiation (1-way ANOVA, Tukey *post hoc,* n=3). Relative expression instead of fold-change versus control was used when no expression was detected in control samples. **D)** Representative immunofluorescence images of differentiated organoids towards the endocrine lineage. Insulin (red) and Neurog3 (green) expression in control and differentiated organoids. White arrowheads indicate insulin^+^ cells. Scalebar=50µm. **E)** Quantification of Neurog3^+^ cells per organoid in control and differentiated organoids (1-way ANOVA, Tukey *post hoc*) and **F)** Insulin^+^ cells per organoid in control and differentiated organoids (Kurskal-Wallis test, Dunn *post hoc*).

Embryonic bipotent progenitors express many ductal cell markers, and progressively lose their progenitor capacity as they mature into ducts^10^. Thus, we analyzed if organoids from different ductal compartments were transcriptionally similar to a bipotent progenitor **(Supplementary_Figure 2C)**. We observed that organoids derived from bigger ducts were closer to a bipotent progenitor based on reduced expression of *Nr5a2, Gata4,* and *Pdx1* while maintaining *Gata6* and *Onecut1* **(Figure 2B)**. We therefore asked if the higher organoid formation capacity of bigger ducts, together with the expression of markers closely related to a bipotent progenitor, could also indicate an increased capacity for endocrine differentiation. Thus, we developed a differentiation protocol that mimics embryonic signaling cues **(Supplementary_Figure 3D).** We observed that medium-big duct-derived organoids display higher capacity for differentiation into endocrine progenitors, showing increased *Neurog3* and *NeuroD1* mRNA levels **(Figure 2C and Supplementary_Figure 3A)** as well as increased Neurog3 protein levels **(Figure 2D-E).** Importantly, organoids derived from bigger ducts also display a higher capacity of differentiation into endocrine lineages, including increased mRNA levels of *Sst* and *Ins* but no *Gcg* or *Ppy* **(Figure 2C and Supplementary_Figure 3B).** Interestingly Insulin^+^ cells did not co-express Neurog3, suggesting that organoid differentiation mimics *in vivo* embryonic development **(Figure 2D-F).** Low levels of acinar markers were detected in all organoids **(Figure 2C).** Finally, expression of ductal markers, although reduced, was still observed in all organoids **(Supplementary_Figure 3C)**. Of note, all experiments with organoids were made with isolated cells to eliminate duct-resident endocrine cells as the contamination could lead to misinterpretation of the results **(Supplementary_Figure 3D).**

**Figure 3:**
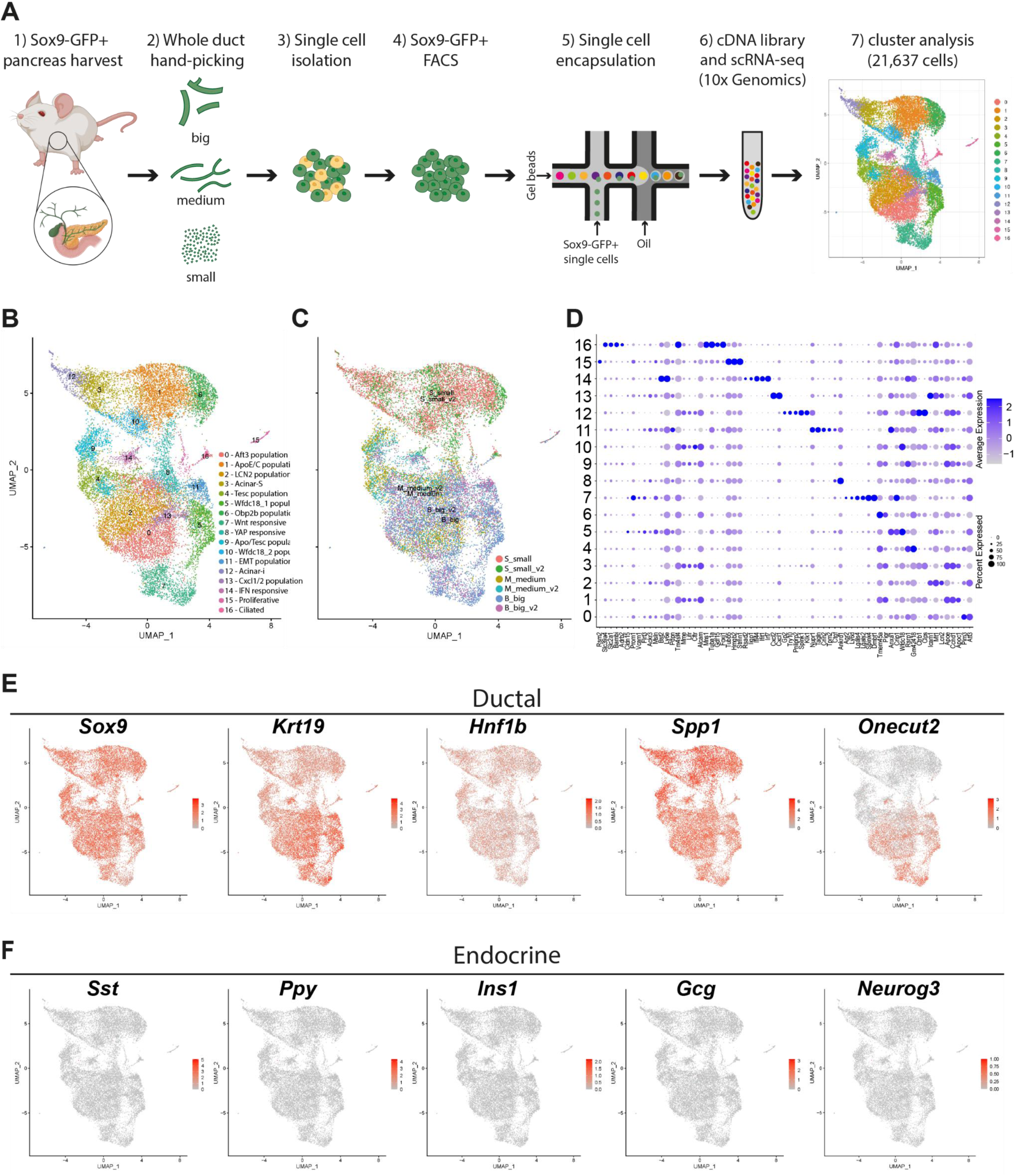
scRNA-seq identifies 15 subtypes of ductal cells. **A)** Schematic representation of the experimental workflow for scRNA-seq **B)** UMAP of 21,637 Sox9:eGFP^+^ is coloured following clustering analysis. **C)** UMAP with samples clustered and coloured by the size of the ducts and experimental replicates. **D)** Dot plot of cell-type-enriched genes per clusters. **E)** UMAP plots showing the expression of pancreatic ductal cell markers. **F)** UMAP plot showing expression of pancreatic endocrine cells: α-cells (*Gcg*), β-cells (*Ins1*), δ-cells (*Sst*), PP-cells (*Ppy*), and endocrine progenitors (*Neurog3*).

### scRNA-seq of ductal cells from mouse adult pancreas identifies novel ductal populations

Encouraged by the observation that ductal compartments behaved differently in organoid cultures, pointing out to the existence of different ductal populations, we comprehensibly characterized ductal heterogeneity by scRNA-seq of isolated Sox9:eGFP^+^ cells from each ductal compartment, excluding the intrapancreatic bile and common biliary duct, followed by FACS. Since each sample was run independently, we were able to later identify the origin of each newly identified cluster **(Figure 3A-C).**

To identify previously unknown ductal populations, we integrated all cells sequenced and filter out doublets and low-quality cells. Our dataset contained 21,637 cells and cluster analysis identified 17 populations with an average of 7,133 UMI/cell and 2,706 genes/cell **(Figure 3B-D and Supplementary_Figure 4A-B).** Clustering analysis showed that cells from small ducts cluster separately **(Figure 3C).** Moreover, we could attribute four clusters derived from small duct fractions (cluster 1, 3, 10, 12), one cluster derived from medium ducts (cluster 4), four clusters derived from big ducts (cluster 2, 5, 7, 13), three clusters derived from medium-big ducts (cluster 0, 9, 11) and four clusters containing mixed cells (8, 14, 15,16) **(Supplementary_Table 1).**

**Figure 4:**
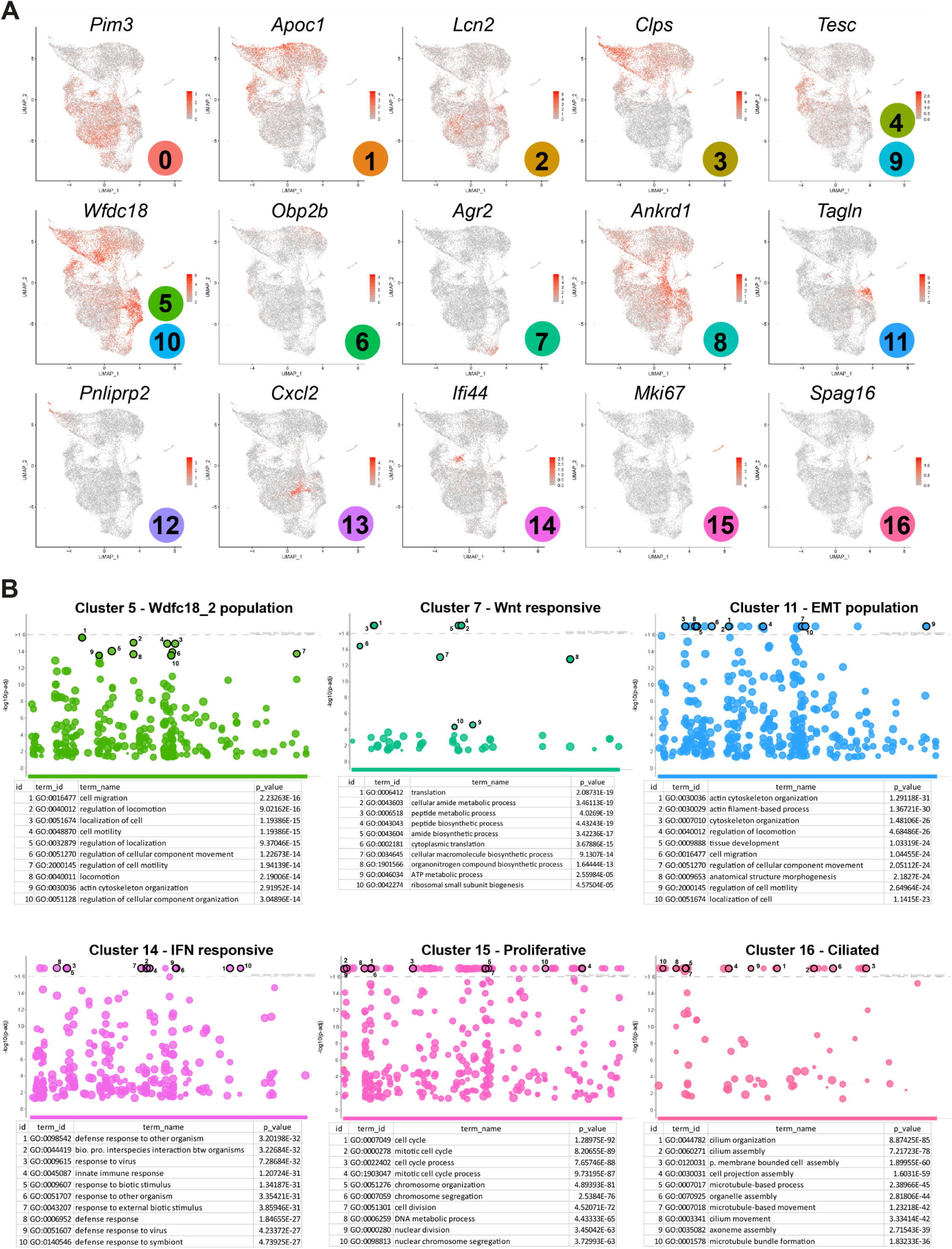
Cluster specific markers and Gene Ontology (GO) analysis of novel ductal populations. **A)** UMAP plots of cluster specific markers for each identified ductal pancreatic population. **B)** Plots showing GO analysis for clusters 5, 7, 11, 14, 15 and 16, with the top 10 significantly gene ontology terms identified.

All clusters expressed ductal markers **(Figure 3E)**, although we observed that *Krt19* expression was lower in clusters mainly derived from small ducts, and *Sox9* and *Spp1* expression was lower in clusters derived from bigger ducts, especially clusters 7 and 5. *Onecut2* expression was absent in small-derived clusters while it was highly expressed in medium-big-derived clusters. Finally, *Hnf1β* expression was evenly distributed **(Figure 3E and Supplementary_Figure 4C).** These results correlate with the qPCR analysis in organoids derived from different ductal fractions **(Figure 2A).**

### Identification of acinar contaminants but not endocrine cells

Further analysis of endocrine lineage-specific markers (*Neurog3*, *Ins*, *Gcg*, *Sst,* and *Ppy*) to investigate the existence of a ductal-endocrine population showed no expression of endocrine markers **(Figure 3F).**

On the other hand, the analysis of acinar-specific markers showed high expression of those in clusters 12 and 3 **(Supplementary_Figure 5A),** suggesting a possible contamination of acinar cells, ambient RNA or early acinar-to-ductal metaplasia (ADM). Surprisingly, cluster 12 and 3 also expressed ductal markers **(Figure 3E).** Thus, to clarify if these clusters represent acinar contaminants or ADM, we sorted Side Scatter^high^ (SSC)/Sox9:eGFP^+^/EpCAM^+^ cells (representing GFP^+^ acinar cells present in our scRNA-seq) and SSC^low^/Sox9:eGFP^+^/ EpCAM^+^ cells (ductal cells) **(Supplementary_Figure 5B).** Staining for acinar (Amylase) and ductal markers (GFP) of the sorted populations clearly showed that SSC^high^GFP^+^ cells were acinar contaminants as most of the amylase^+^ cells were GFP^-^ but show a GFP^+^ CAC attached **(Supplementary_Figure 5C)**.

**Figure 5:**
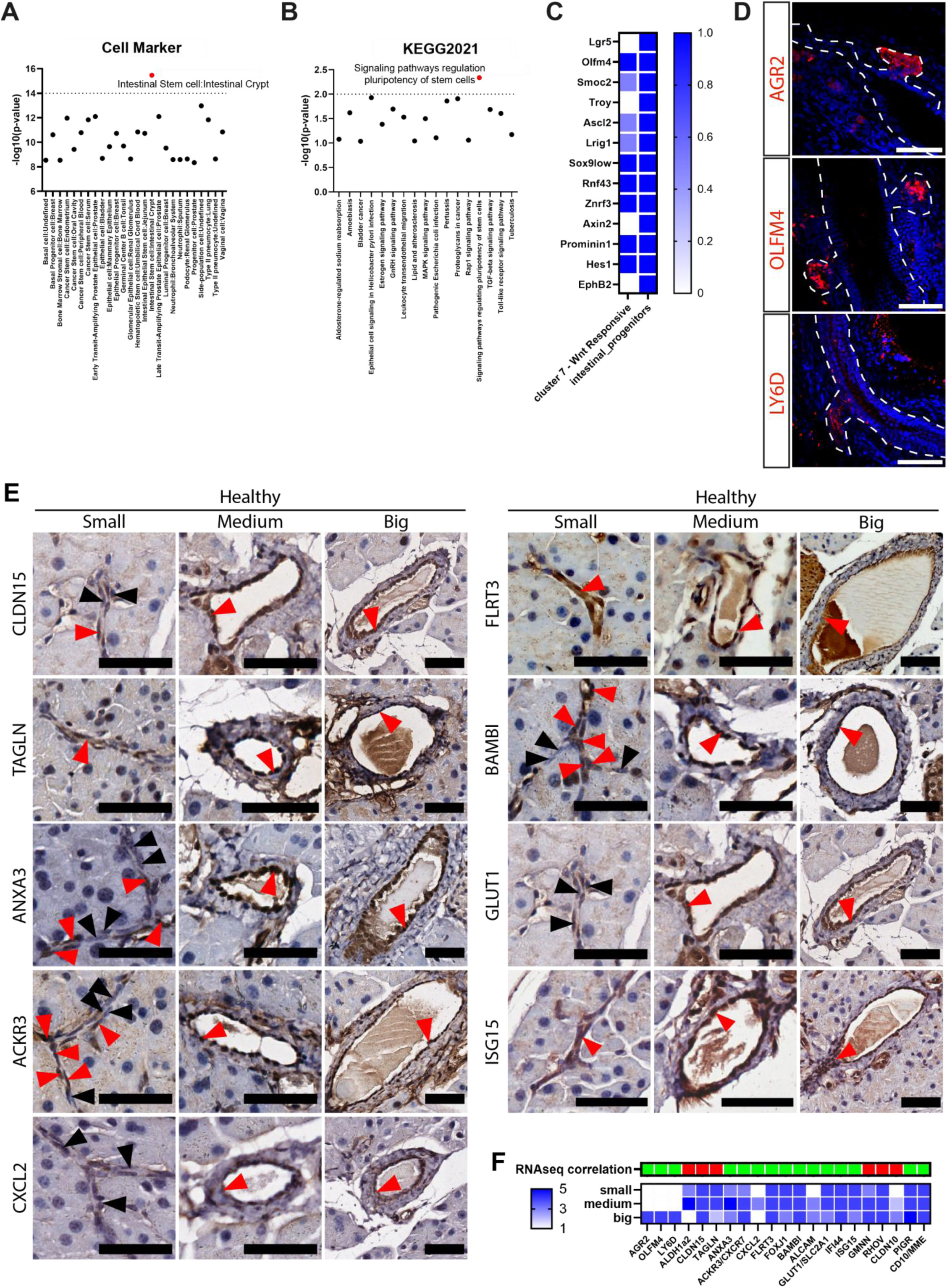
Detection of novel ductal markers in healthy pancreatic tissue, including Wnt-responsive-population in pancreatic ductal glands. **A)** Plots showing Enrichr analysis^55^ for Wnt-responsive-population in CellMarker Augmented (hits=-log(p-value)>8) and **B)** KEGG2021 (hits=-log(p-value)>1). **C)** Gene expression comparison between Wnt-responsive-population and intestinal progenitors^35^. **D)** Representative wholemount staining of the main pancreatic duct (white dashes) for Wnt-responsive-populations specific markers, AGR2, OLFM4 and LY6D (red) and nuclei (DAPI, blue), showing positive PDGs. Scalebar=300µm. **E)** Representative images of immunohistochemistry detection of CLDN15, TAGLN and ANXA3 (EMT-population), ACKR3 (YAP-responsive-population), CXCL2 (Cxcl1/2-population), FLRT3 (medium-big ducts), BAMBI (ciliated-population), GLUT1 (expression in some ductal cells) and ISG15 (IFN-responsive-population) in healthy pancreas (red arrows indicating positive cells, black arrow negative cells). Scalebar=50µm. **F)** Heatmap plot shows qualitative correlation (green) or no correlation (red) between protein expression and scRNA-seq data. Heatmap plot shows, in blue, qualitative analysis of marker expression in small, medium, and big ducts in healthy tissue.

Furthermore, although both acinar clusters expressed acinar markers, these were highly expressed in cluster 12, indicating a more stable secretory phenotype **(Supplementary_Figure 5D).** Recent snRNA-seq studies^19^ described two acinar populations: acinar-i (idling-acinar) and acinar-s (secretory-acinar). In our scRNA-seq and immunofluorescence, we identified Cluster 12 as acinar-s-population (Amylase^high^) and Cluster 3 as acinar-i-population (Amylase^low^) **(Supplementary_Figure 5D-E)**.

### Novel ductal populations

Significantly differentially expressed genes when comparing a cluster to all other clusters highlighted the existence of previously unidentified ductal populations **(Figure 3B-D**, **Figure 4A and Supplementary_Table 2).**

Interestingly, clusters derived from small ducts allowed for the identification of genes uniquely or highly expressed in CACs/TD. Cluster 6 (Obp2b-population) is characterized by the expression of *Obp2b*, known to be expressed in the mammary gland^25^ but not in the pancreas. In addition, Cluster 1 (ApoE/C-population), expresses apolipoproteins (*ApoC1*/*ApoE*) that have been associated with PDAC prognosis^26^. Cluster 1 also displays high levels of *Mme* and *Cldn10*. Mme is highly expressed in neuroendocrine tumors^27^, and Cldn10 is already known to be expressed in TD^28^. Cluster 10 (Wfdc18_2-population), is characterized by high expression of *Wfdc18* that has been related to PDAC^29^. Interestingly, our data indicates that *Prox1* expression is higher in CACs/TD. Prox1 plays an important role in pancreas morphogenesis. It is expressed in pancreatic progenitors, and as development proceeds, its expression gets restricted to islets and some ductal cells^30^. Finally, *Cftr* is highly expressed in small ducts-derived/containing clusters, and mutations in *Cftr* produce cystic fibrosis^31^. Therefore, our data suggests that CACs/TD could be the main players in the exocrine damage/blockage observed in cystic fibrosis patients.

Cluster 4 (Tesc-population), derived from medium size ducts, is characterized by the expression of *Tescalcin* that has recently been reported as a regulator of cancer progression^32^.

Clusters derived from big ducts are characterized by the expression of previously unidentified ductal markers. Cluster 2 (Lcn2-population) is characterized by the expression of *Lcn2* and *Icam1*. Increased expression of both has been observed in pancreatitis, PanINs, and PDAC^33,34^. Cluster 5 (Wfdc18_1-population), is characterized by high expression of *Wfdc18* like Cluster 10. Interestingly, Cluster 7 (Wnt-responsive-population) expresses many markers shared with progenitors in other tissues, including *Olfm4*, *Ly6D*, *Agr2* and *Hes1*, therefore displaying a progenitor-like transcriptome similar to intestine^35^ and prostate^36^, including Wnt-responsive genes (*Ascl2*, *Rnf43* and *Znrf3*). Cluster 13 (Cxcl1/2-population), is characterized by high expression of *Cxcl1* and *Cxcl2*. Their expression in iCAFs correlates with an increase invasive phenotype in PDAC^37^.

Several clusters were derived from medium-big ducts, Cluster 0 (Atf3-population) expresses high levels of *Atf3*. *Atf3* deletion in acinar cells decreased pancreatitis-induced ADM, PanIN formation and PDAC^38^. Cluster 9 (Apo/Tesc-population), shares most of the markers with Clusters 4 or 1. Cluster 11 (EMT-population) is characterized by the expression of *Tagln*, *Tpm2*, and *Tpm1* that have been identified as markers of myCAFs in PDAC^39^, although these cells maintain the expression of ductal cells markers, thus suggestive of an epithelial-mesenchymal transition state.

Finally, among the clusters composed of mixed compartments, Cluster 8 (YAP-responsive-population) is highly enriched in YAP target genes, including *Ankrd1*, *Ctgf* and *Cyr61*. Hippo-signaling has been found to play key roles in progenitor maintenance, renewal, proliferation, and differentiation in pancreatic progenitors^40^. Unexpectedly, Cluster 14 (IFN-responsive-population) is highly enriched in antigen-presenting and interferon response genes (*Isg15*, *Ifitm3, Ifi17l2a* and *Irf7)* that have recently been related to a ductal population enriched in islets of T1D patients that could play a role in immune eviction^41^. On the other hand, Cluster 15 (proliferative-population), represents a subset of cycling ductal cells. Finally, Cluster 16 (ciliated-population) is characterized by high expression of cilia markers, specifically motile cilia, such as *SPAG16* and 17, and *Foxj1*.

### Wnt-responsive-population is located in pancreatic ductal glands

Gene Ontology analysis showed many biological processes shared by several clusters, such as tissue development, positive regulation of cellular processes, and cellular respiration **(Supplementary_Figure 6 and Supplementary_Table 3).** Remarkably some clusters display unique terms, such as, Cluster 14, IFN-responsive-population, enriched in immune response and cellular response to interferon pathways. Clusters 5 and 11, Wfdc18_1-population and EMT-population respectively, were enriched in terms related to cell migration and differentiation. Cluster 15, proliferative-population, was enriched in mitotic processes and RNA splicing and export. Cluster 16, ciliated-population, was enriched in cilium assembling, docking, and transport **(Figure 4B and Supplementary_Figure 6).** Thus, our scRNA-seq results show ductal heterogeneity and highlight previously unidentified ductal populations that could play different roles in pancreatic pathogenesis and regeneration.

**Figure 6:**
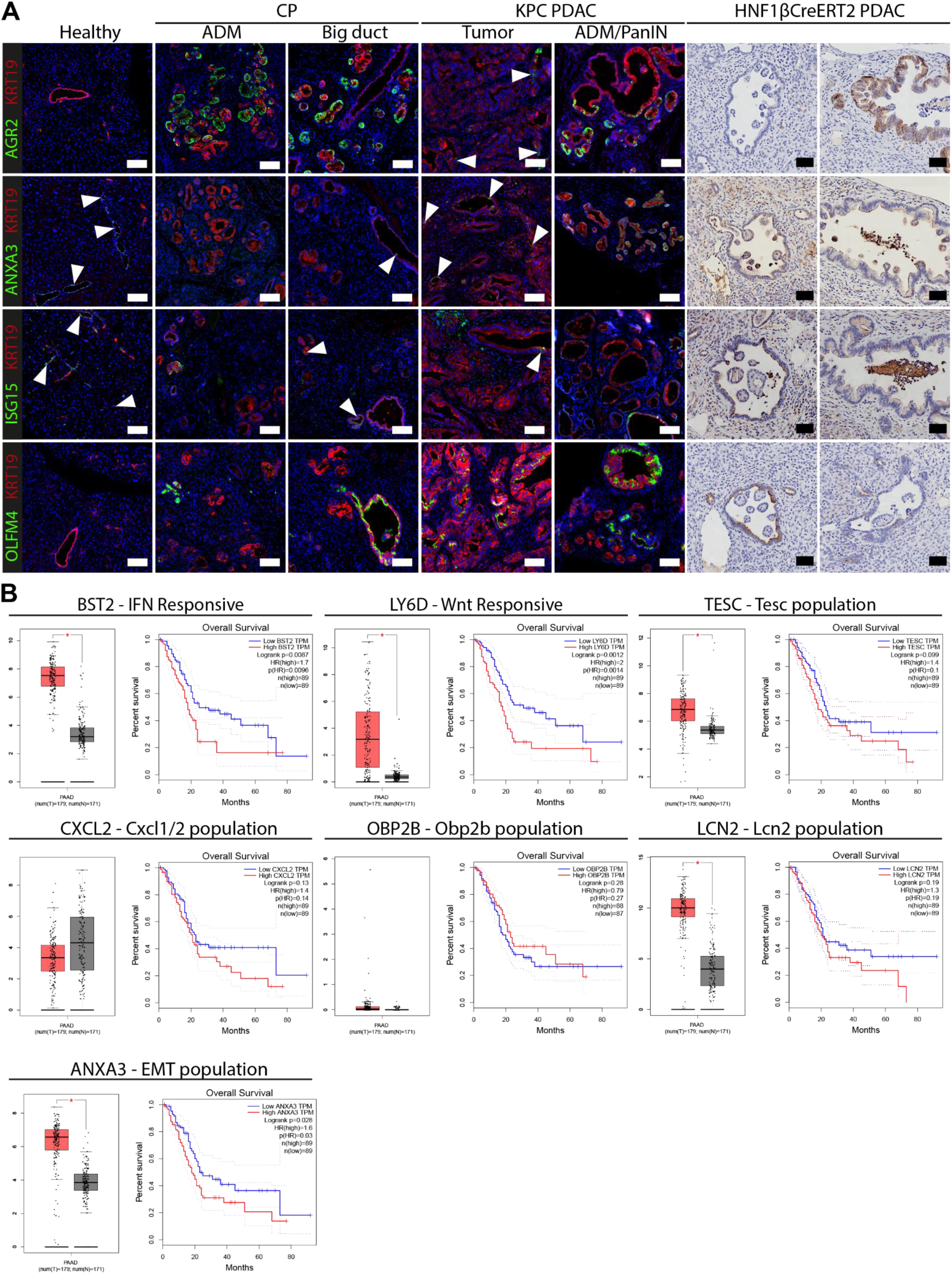
Characterization of ductal populations in adult pancreas, chronic pancreatitis and PDAC. **A)**Representative immunofluorescence/immunohistochemistry images of healthy, chronic pancreatitis and PDAC samples from KPC (LSL-KrasG12D/+;LSL-Trp53R172H/+;Pdx-1-Cre) and HPC (*LSL-Kras^G12V^;*Trp53R172H/+;Hnf1b^CreERT2^) mice. Staining of population markers, AGR2 and OLFM4 (Wnt-responsive-population), ISG15 (INF-responsive-population) and ANXA3 (EMT-population) were detected in green and pan-ductal marker KRT19 in red. Representative images of ADM, tumor and PanINs are shown per every marker analyzed. Arrowheads indicate coexpression of population markers and KRT19. Scalebar=50µm. **B)** mRNA expression of population markers BST2 (IFN-responsive-population), LY6D (Wnt-responsive-population), TESC (Tesc-population), CXCL2 (Cxcl1/2-population), OBP2B (Obp2b-population), LCN2 (Lcn2-population) and ANXA3 (EMT-population) in human PDAC samples and healthy pancreas via GEPIA (http://gepia.cancer-pku.cn/) database including TCGA and GTEX data. Box plot represents tumor (T) in red (179 samples) and normal tissue (N) in gray (171 samples). X-axis represents mRNA expression as log2(TPM+1). Significant differences are shown (*α<0.001). Kaplan-Meier survival curves are plotted using GEPIA where median group survival cutoff of 50% is showed, as well as Hazards Ratio and 95% confidence intervals.

Interestingly, Wnt-responsive-population was enriched in biological processes related to ribosome biogenesis and translation **(Figure 4B),** which are linked to stemness, especially in quiescent stem cells^42^. When analyzing KEGG pathways with genes overexpressed ≥1.5-fold in this population, we observed an enrichment of signaling pathway regulating pluripotency of stem cells **(Figure 5A and Supplementary_Table 4),** and CellMarker analysis^43^ showed high similarities with intestinal and basal stem cells and side population **(Figure 5B and Supplementary_Table 4),** thus suggesting a stem-like phenotype of this populations. Many of the classical markers expressed by Lgr5 intestinal stem cells, but not Lgr5 **(Supplementary_Figure 3E)**, were expressed by Wnt-responsive-population **(Figure 5C).** This population also expresses markers previously identified in extrahepatic biliary cells of the common biliary duct, like Dmbt1 and Ly6D^44^. We manually dissected out the intrapancreatic bile and common biliary duct for our scRNA-seq experiments, therefore these results suggested that Wnt-responsive-population is like a population located in the extrahepatic biliary duct. Surprisingly, staining for Wnt-responsive-population markers (Ly6D, Agr2, or Olfm4), showed its expression in budding/outpouching structures in the main duct, named pancreatic ductal glands (PDGs)^45^ **(Figure 5D)**. Interestingly, PDGs have been suggested to be niche compartments both in mouse and human pancreas, displaying progenitor capacities and playing a role in regeneration of exocrine pathologies^45,46^.

### Characterization of ductal populations in adult pancreatic tissue and injury models

Besides Wnt-responding-population, we characterized the expression of other populations’ specific markers by immunohistochemistry in healthy pancreatic tissue **(Figure 5E and Supplementary_Figure 7).** Significantly, we confirmed protein expression of most of the cluster-specific markers: Bst2 and Isg15 for IFN responsive-population, Tagln and AnxA3 for EMT-population, and Foxj1 for ciliated-population. Flrt3 for medium/big-derived-populations and Aldh1a2 for Wfdc18_1-population **(Figure 5E)**. We confirmed the enrichment correlation of markers in the studied ductal compartments and our scRNA-seq data for 71.4% (15 out of 21) **(Figure 5F)**. Next, we characterized the expression of specific markers in chronic pancreatitis (CP) and observed that Olfm4, Agr2, and Isg15 are heterogeneously expressed in acinar to ductal metaplasia regions (ADM) in CP samples, thus suggesting a putative contribution to pancreatitis pathogenesis of the Wnt-responsive-population and IFN-responsive-population **(Figure 6A).**

**Figure 7:**
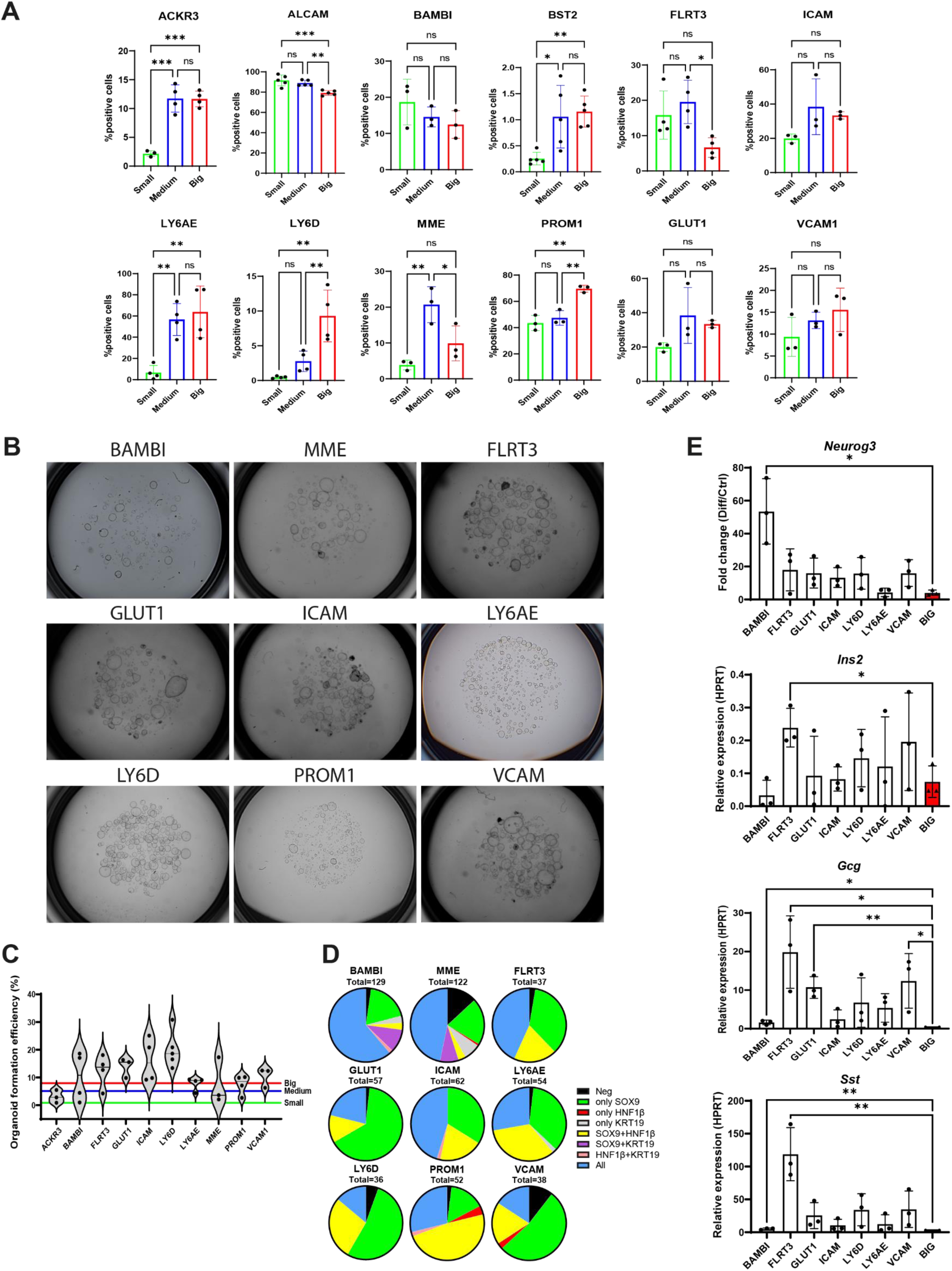
Isolation of newly identified populations and *in vitro* characterization in organoid cultures upon endocrine differentiation. **A)** Bar plots showing quantification of ductal populations distribution in small, medium, and big-enriched ductal fractions by FACS (1-way ANOVA, Tukey’s *post hoc*, n=3-5). **B)** Representative images of organoid formation efficiency upon 10-days in culture. **C)** Violin plot showing quantification of organoid formation efficiency (n=3-5). Population mean is shown per violin plot. Colored lines indicate the mean of organoid formation in big (red), medium (blue) and small (green) duct-derived organoids. **D)** Pie charts showing quantification of ductal markers expression by antibody staining (SOX9, HNF1β and KRT19) in population-derived organoids. **E)** qPCR analysis on population-derived organoids upon differentiation for *Neurog3*, *Ins2*, *Gcg* and *Sst* (Student t-test vs big (red), n=3).

In the last years, several laboratories have described acinar and ductal cells as origins of PDAC^4,22,47,48^. Ductal-derived PDACs, characterized by the expression of Agr2^49^ or IFN signaling-related genes^47^, are more aggressive and have a poorer outcome. Interestingly, Wnt-responsive-population is characterized by the expression of Agr2 and IFN-responsive-population by IFN-related genes, such as ISG15 and Bst2. Thus, we characterized the expression of those markers in mouse PDAC samples, either in tumors where Kras mutation and P53 deletion is induced by the Pdx1-Cre (progenitor-derived, K-rasLSL.G12D/+;p53R172H/+;Pdx1Cre^50^) or by the HNF1bCreERT2 (ductal-derived, K-rasLSL.G12D/+;p53R172H/+;HNF1bCreERT2 or K-rasLSL.G12V/+;HNF1bCreERT2^22,48^) system. In Pdx1-derived tumors, we observed scattered expression of Agr2, Olfm4, AnxA3, and Isg15 in the tumor and ADM and PanIN lesions. In ductal-derived tumors, we observed two different patterns in PanINs, uniformly expressing PanINs and PanINs lacking marker expression **(Figure 6B)**. Interestingly, specific markers from Wnt-responsive, IFN-responsive and EMT-populations had higher outcome predictive values in human samples than genes in other populations such as Obp2b-population, Cxcl1/2-population, Lcn2-population or Tesc-population **(Figure 6B).** Our results suggest the possibility of different roles of novel populations in the pathophysiology of exocrine pancreatic diseases, as well as different ductal cell of origin of PDAC that require further investigation to increase our understanding of the biology and PDAC heterogeneity.

### Isolation of newly identified populations and *in vitro* characterization of exocrine/endocrine differentiation capacity

To further study newly defined populations in organoid cultures we intersected our scRNA-seq data with the human protein atlas data on membrane proteins^51^ and identified antibodies against extracellular epitopes of these proteins. Isolation of ductal compartments followed by antibody staining of specific surface markers before FACS confirmed the enrichment of the ductal populations in the compartments identified by scRNA-seq in 10 out of the 15 markers analyzed **(Figure 7A, Supplementary_Figure 8A)**. We isolated Ly6D^+^ cells (Wnt-responsive-population), Bambi^+^ cells (ciliated-population), Bst2^+^ and Ly6A/E^+^ cells (INF-responsive-population and cells from medium/big ducts), Vcam1^+^ cells (Wfdc18_1-population), Icam1^+^ cells (Atf3, ApoE/C, and Cxcl1/2-populations). Ackr3^+^ and Flrt3^+^ cells (medium-big duct-derived clusters) and CD166/Alcam^+^ cells (pan-ductal). CD133/Prominin, was selected as stem cell marker already used to test ductal cell stemness (reviewed^52^). Glut1/Slc2a1 was used to isolate a fraction of ductal cells, although it does not define a cluster. Finally, Mme was used to enrich for small ductal cells, although the expression pattern observed by FACS did not correlate with scRNA-seq expression data **(Figure 7A and Supplementary_Figure 8A)**.

The organoid formation capacity of the isolated populations based on Sox9:eGFP expression and specific surface markers was interrogated by plating the same cell density **(Figure 7B-C).** All ductal isolated populations formed organoids although at different efficiency. The populations with higher organoid formation efficiency were derived from medium-big ducts including: Ly6D (20.05%±2.25), Icam1 (16.24%±7.98), Glut1 (13.89%±3.55), Vcam1 (10.38%±3.55), Ly6A/E (7.95%±2.34), Flrt3 (12.43%±5.88), and Bambi (10.28%±8.87). Some of the isolated populations display low organoid formation capacity below 7.93% observed in big ducts **(Figure 1J)**, including Prominin, Ackr3 and Mme (7.43%±3.48, 3.04%±2.3 and 7.65%±8.39, respectively). We also quantified organoid size **(Supplementary_Figure 8B),** showing different growth capacities. Notably, organoid heterogeneity, based on ductal markers, was highly reduced **(Figure 7D).** Only populations with an organoid formation efficiency higher than big ducts were further analyzed.

*In vitro* differentiation of the isolated populations into the endocrine lineages showed that Bambi^+^ cells display a significant capacity to differentiate into endocrine progenitors, up to 50 times fold-increase of *Neurog3* expression versus control organoids (53.49±19.89) **(Figure 9E**). Other isolated populations, Flrt3, Glut1, Icam1, Ly6D and Vcam display increased capacity for differentiation into *Neurog3* expressing cells when compared to big-duct derived organoids, although they do not reach statistical significance. Surprisingly, Bambi-derived organoids did not display high capacity of differentiation into endocrine lineages based on *Ins*, *Gcg* and *Sst* mRNA expression, instead Flrt3-derived organoids display higher capacity of differentiation into all endocrine linages **(Figure 9E)**. Curiously, Ly6D-derived organoids show higher capacity to differentiate into acinar cells based on the expression of digestive enzymes, such as amylase **(Supplementary_Figure 8C)**.

## DISCUSSION

Pancreatic heterogeneity, at single-cell resolution, has been mainly exploited to study pancreatic endocrine cells, thus ductal heterogeneity has not been deeply studied either in mice or humans. The first glimpse of such heterogeneity was highlighted in recent publications^17,18^, although these studies do not include cells from all ductal compartments and only a couple of thousand cells were analyzed. We have now generated, for the first time, a murine pancreatic ductal cell atlas of the entire ductal epithelium from the tip (CACs) up to the main duct including PDGs. Our results have uncovered an unprecedented level of heterogeneity in the ductal compartment. Interestingly, recent work Fasolino *et al.*^41^ discovered a subset of ductal cells that acquires a signature of tolerogenic dendritic cells in an attempt at immune suppression in T1D donors, highlighting a novel role of ductal cells in T1D pathogenesis. Interestingly, our scRNA-Seq analysis demonstrated the presence of this population scattered through the ductal epithelium. Thus, further investigations are needed to understand the role of this population in the development and pathophysiology of diabetes or exocrine diseases.

We have defined an *in vitro* protocol to efficiently induce differentiation of organoids into Neurog3^+^ progenitors and to a lesser extent into hormone-producing cells. Interestingly, bigger ducts and derived organoids express lower levels of *Spp1*, thus our results are in concordance with the recent observations demonstrating that inhibition of Spp1 in ductal cells induces insulin expression *in vitro*^18^. Our scRNA-seq data demonstrates the coexistence of 15 ductal populations, some of which were not previously identified^17,18^ and demonstrated functional differences within the ductal epithelia. scRNA-seq data did not contain endocrine cells or Neurog3^+^ cells, as suggested by Gribben *et al.*^13,53^. The absence of Neurog3^+^ cells in our study is most likely due to the selection of Sox9:eGFP^+^. The likelihood of the presence of a pool of dedicated progenitors in the adult pancreas is low, thus β-cell neogenesis has been controversial. Most of the lineage tracing studies, using pan-ductal-specific Cre-lines^12^, did not find a quantifiable and physiologically relevant contribution of β-cell neogenesis from ducts. These previous results do not disprove the plasticity of ductal cells *in vitro*, in organoid cultures, as we have demonstrated here showing that different ductal compartments behave differently in organoid cultures at different levels, from organoid formation to endocrine/exocrine differentiation.

We further showed that upon cell sorting isolation of novel populations derived organoids Bambi, Flrt3, Glut1, Icam1, Ly6D and Vcam display over 10% organoid formation efficiency, although organoid formation did not correlate with an increased differentiation capacity into endocrine differentiation potential, at least under the same media conditions. Surprisingly, organoids derived from the ciliated-population, based on Bambi expression, display high differentiation capacity towards endocrine progenitors but no hormone-producing cells. Intriguingly, Flrt3-derived organoids show high efficiency of differentiation into hormone-expressing cells. Interestingly, Flrt3 has been found to be expressed in pancreatic multipotent progenitors at E9.5^54^. Strikingly, Wnt-responsive-population located in PDGs displays stemness features but does not display high endocrine differentiation capacity, instead it shows high exocrine regenerative capacity. On the other hand, future manipulations to promote β-cell differentiation from organoids are needed to develop efficient protocols. We have used a standardized media for organoid formation and differentiation, thus further investigation of the signaling cues required to differentiate novel ductal populations into the endocrine lineage may differ between them. Nevertheless, our results demonstrated a putative role of novel ductal populations in endocrine and exocrine regeneration.

We demonstrate that the newly identified populations might play a role in pancreatic exocrine pathogenesis as its expression increases in ADM regions in CP. In addition, recent reports highlighted acinar and ductal cells of origin of PDAC. Interestingly, our results highlighted the existence of three ductal populations, Wnt-responsive, IFN-responsive and EMT-population showing higher outcome predictive values in human PDAC samples than genes in other populations. Thus, our data strongly suggests the possibility of the existence of more than one ductal cell of origin of PDAC. If uncovered, it would be key to investigate the differences in tumor development and biomarkers according to a ductal cell origin to design a personalized medicine strategy.

Considering our discovery of previously unidentified ductal populations, we unmask the potential roles of specific ductal populations in endocrine/exocrine regeneration and exocrine pathogenesis. Therefore, our results compel the need to reinterpret the cellular pathogenesis of pancreatic diseases and to revisit previous lineage tracing experiments using pan-ductal markers by generating new tracing tools, to study ductal-specific populations to investigate its regenerative potential and its role in exocrine pathogenesis *in vivo*.

## AUTHOR CONTRIBUTIONS

M.R. conceived, supervised, and coordinated the study. A.F performed organoid cultures and qPCR analysis. A.F, A.H and J.C performed flow cytometry and sorting experiments.

J.C. performed immunohistochemistry and imaging on healthy pancreas and PDAC samples and imaging of organoids, as well as manuscript figures. A.H. performed cytospin experiments and acinar cell isolation. I.R and K.C performed immunofluorescence and imaging acquisition of injury models and tumor samples. L.M. performed gene ontology analysis. L.P. supervised A.F and corrected the manuscript. J.M.B-L performed the experiments to generate ductal derived tumors. Y.J.W performed single-cell computational analysis. M.R. wrote the manuscript with input from all the authors.

## DATA AVAILABILITY

Accession numbers pending, data will be available in an online repository.

## ABBREVIATIONS

3D: Three dimensional
acinar-i: idling acinar
acinar-s: secretory acinar
ADM: Acinar-to-ductal
Agr2: Anterior Gradient 2 Protein Disulphide Isomerase Family Member
ALDH1: Aldehyde Dehydrogenase 1
ALK3: Bone Morphogenetic Protein Receptor Type 1A
Amy2a: amylase
Ankrd1: Ankyrin Repeat Domain 1
AnxA3: Annexin A3
ApoC1: Apolipoprotein C-I
ApoE: Apolipoprotein E
Ascl2: Achaete-Scute Family BHLH Transcription Factor 2
Atf3: Activating Transcription Factor 3
Bambi: BMP And Activin Membrane Bound Inhibitor
Birc5: Baculoviral IAP Repeat Containing 5
Bst2: Bone Marrow Stromal Cell Antigen 2
CAC: Centroacinar cells
CAII: Carbonic Anhydrase 2
CD166/ALCAM: Activated Leukocyte Cell Adhesion Molecule
Cela2a: Chymotrypsin Like Elastase 2A
CFTR: Cystic Fibrosis Transmembrane Conductance Regulator
Cldn10: Claudin 10
Cldn15: claudin 15
CP: chronic pancreatitis
Cpa2: Carboxypeptidase A2
Ctgf or CCN2: connective tissue growth factor
Ctrb1: Chymotrypsinogen B1
Cxcl1: C-X-C Motif Chemokine Ligand 1
Cxcl2: C-X-C Motif Chemokine Ligand 2
Cyr61: Cysteine-rich angiogenic inducer 61
DEG: Significantly differentially expressed genes
Dmbt1: Deleted In Malignant Brain Tumors 1
EMT: Epithelial to Mesenchymal Transition
EpCAM: Epithelial Cell Adhesion Molecule
Foxj1: Forkhead Box J1
Gata4: GATA Binding Protein 4
Gata6: GATA Binding Protein 6
Gcg: Glucagon
GFP: Green fluorescent protein
Ghrl: Ghrelin
Glut1/Slc2a1: Solute Carrier Family 2 Member 1
Gp2: Glycoprotein
Hes1: Hes Family BHLH Transcription Factor 1
HNF1β: HNF1 Homeobox B
Hnf6: Onecut1 or One Cut Homeobox 1
IAD: intracalated ducts
iCAF: inflammatory fibroblasts
Icam1: Intercellular adhesion molecule 1
IED: intercalated ducts
Ifi17l2a: Interferon Alpha Inducible Protein 27 Like 2
Ifitm3: Interferon Induced Transmembrane Protein 3
IFN: Interferon
Ins: Insulin
Irf7: Interferon Regulatory Factor 7
Irf9: Interferon Regulatory Factor 9
Isg15: ISG15 Ubiquitin Like Modifier
Krt19: Cytokeratin 19
Krt7: Cytokeratin 17
Lcn2: Lipocalin2
Lgr5: Leucine Rich Repeat Containing G Protein-Coupled Receptor 5
Ly6a/e: Lymphocyte Antigen 6 Complex, locus A or Stem Cell Antigen-1
Ly6D: Lymphocyte Antigen 6 Family Member D
MD: main duct
MiK67: Marker of Proliferation Ki-67
Mme/CD10: Membrane Metalloendopeptidase
MUC1: Mucin 1, Cell Surface Associated
myCAF: myofibroblastic CAF
Neurog3: Neurogenin 3
Nr5a2: Nuclear Receptor Subfamily 5 Group A Member 2
Obp2b: Odorant-binding protein B
Olfm4: Olfactomedin 4
Onecut2: One Cut Homeobox 2
P2RY1: Purinergic Receptor P2Y1
PanIN: pancreatic intraepithelial neoplasia
PDAC: Pancreatic Ductal Adenocarcinoma
PDG: Pancreatic ductal gland
Pdx1: Pancreatic And Duodenal Homeobox 1
Pigr: Polymeric Immunoglobulin Receptor
Ppy: Pancreatic Polypeptide
Prox1: Prospero Homeobox 1
Prss2: Serine Protease 2
Ptf1a: Pancreas Associated Transcription Factor 1a
RA: retinoic acid
RhoV: Ras Homolog Family Member V
Rnf43: Ring Finger Protein 43
snRNA-seq: Single Nuclei RNA-sequencing
scRNA-seq: Single cell RNA-sequencing
Sox9: SRY-Box Transcription Factor 9
SPAG: Sperm Associated Antigen
Spink: Serine Peptidase Inhibitor Kazal Type 1
Spp1: Secreted Phosphoprotein 1
SSC: Side Scatter
Sst: Somatostatine
Sycn: Syncollin
T1D: Type 1 diabetis
Tagln: Transgelin
TD: terminal ducts
TESC: Tescalcin or calcineurin B homologous protein 3
Tff2: Trefoil Factor 2
TGF-β: Transforming growth factor β
Tmem45a: Transmembrane Protein 45A
Tpm1: Tropomyosin 1
Tpm2: Tropomyosin 2
Vcam1: Vascular Cell Adhesion Molecule 1
Wfdc18: WAP four-disulfide core domain 18
YAP: yes-associated protein 1
Znrf3: Zinc And Ring Finger 3

## ACKNOWLEDGEMENTS

This research was supported by Ministerio de Ciencia e Innovación (MICIN) (SAF2015-73226-JIN, PID2019-106160RB-I00 and RYC-2017-21950) to M.R. A.H. was supported through a fellowship from MICIN (PRE2020-092678). J.B. was supported by NIH grant support number: 1 R01 CA277161-01A1. We thank IDIBELL’s animal facility and mouse unit, especially Antoni Ventura, for housing and maintenance of the Sox9:eGFP line. We thank IDIBELL’s flow cytometry facility, especially J.M. Andrés Vaquero. We thank IDIBELL’s histology facility, especially Lola Mulero and Jose Llamas, and imaging facility, especially Joan Repullès and Saioa Mendizuri. We thank Donald Allen for critical comments and corrections on the manuscript. Drawings created with BioRender.com.

---

**Supplementary Figure 1.**
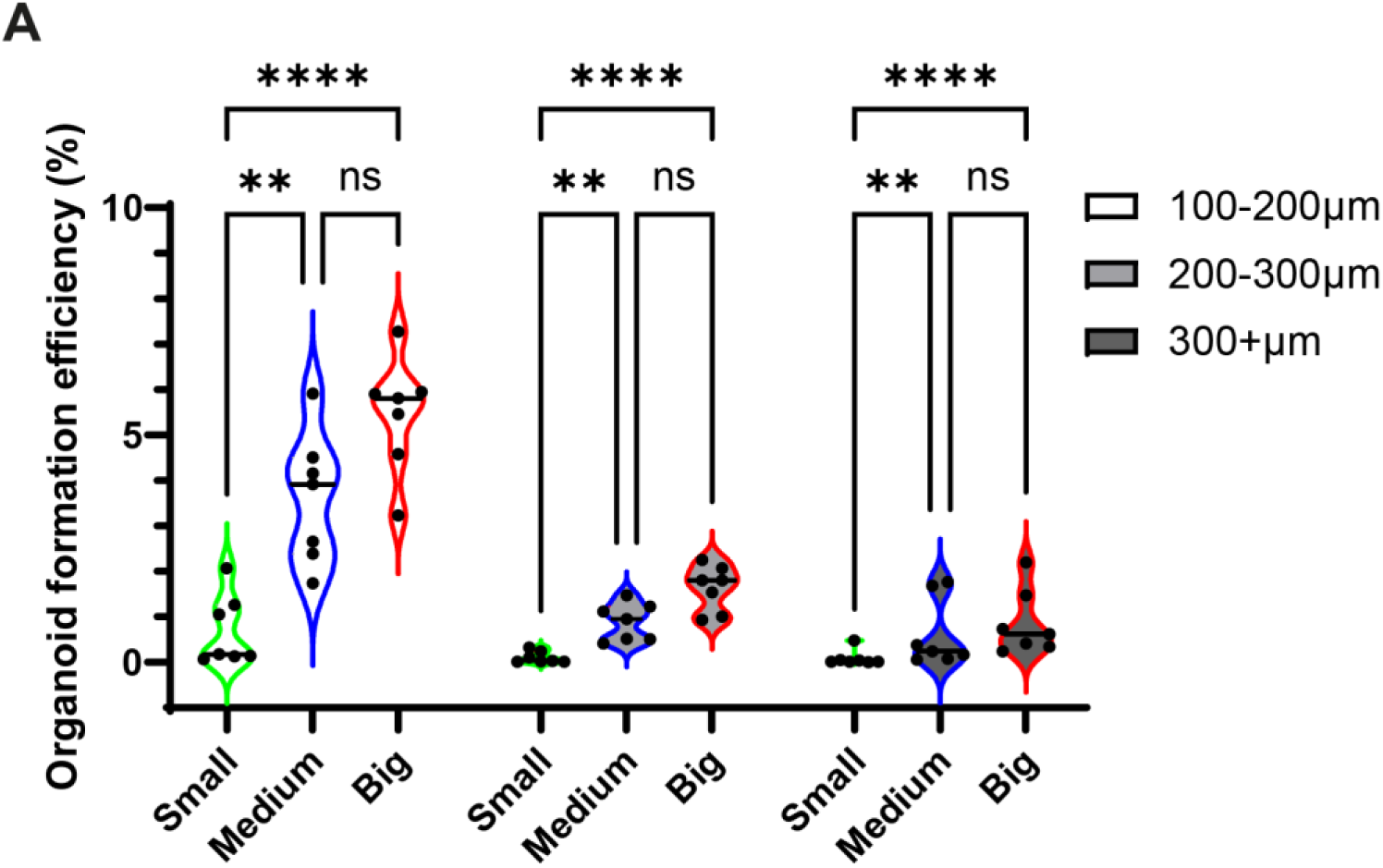

**Supplementary Figure 2.**
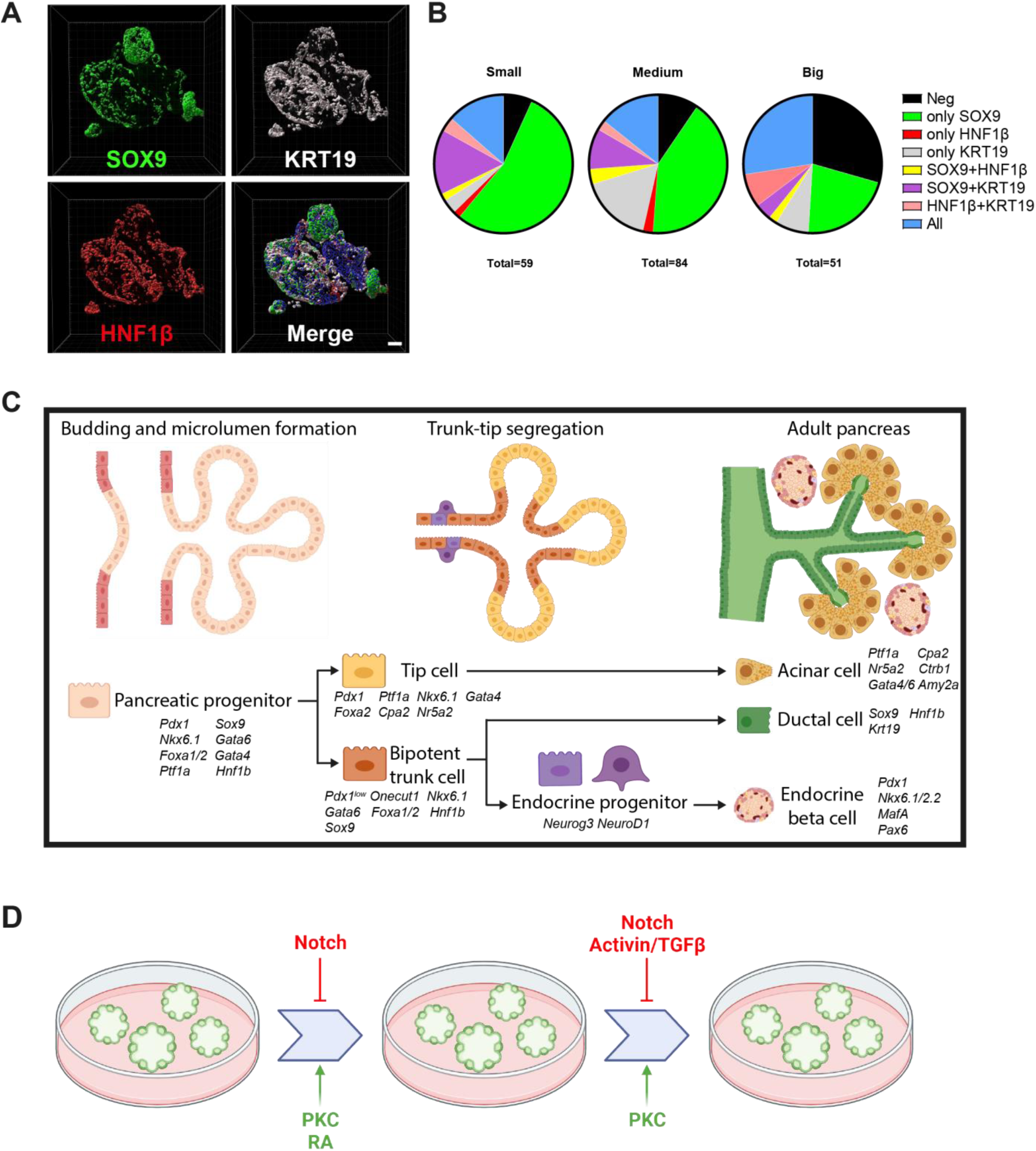

**Supplementary Figure 3.**
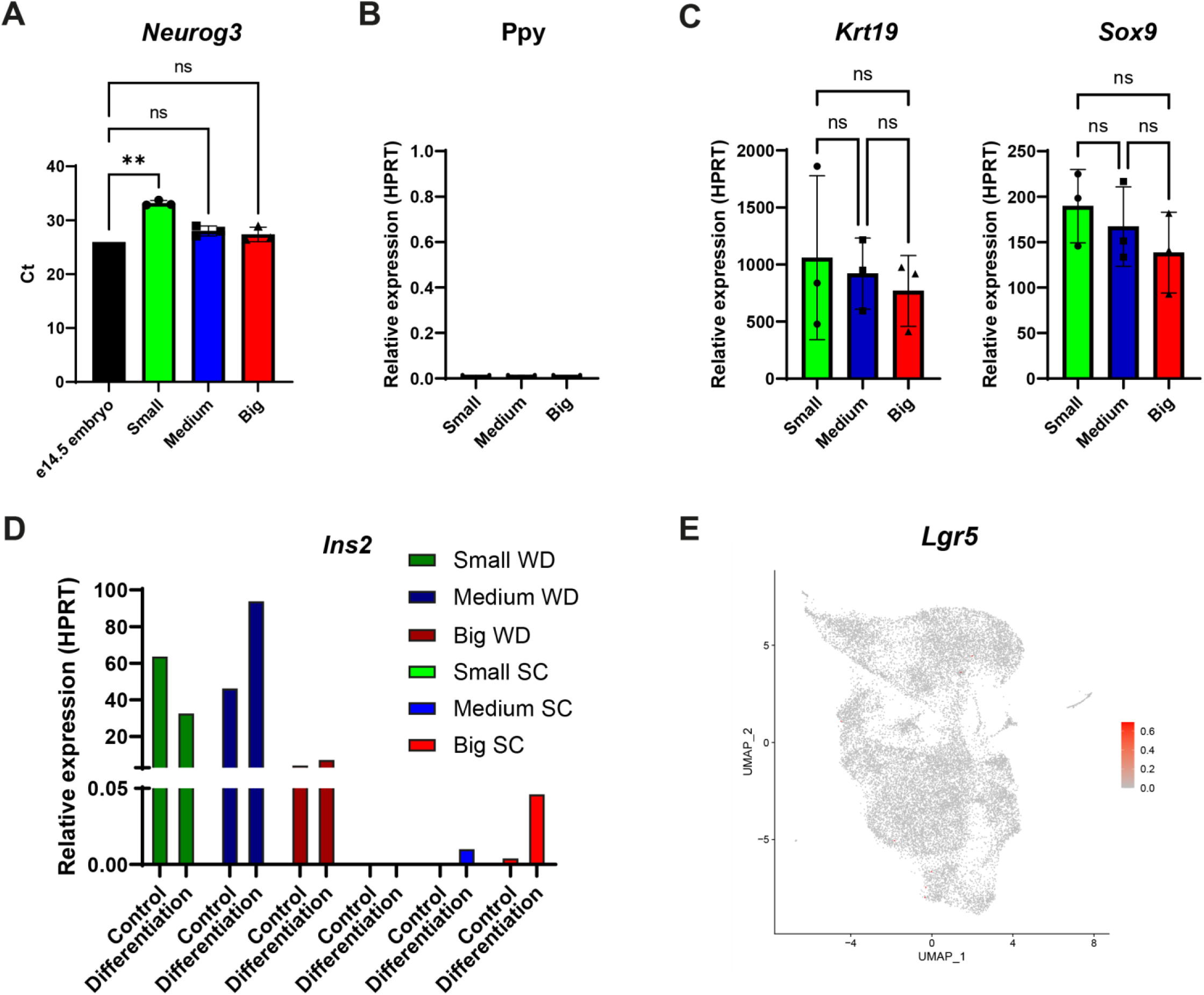

**Supplementary Figure 4.**
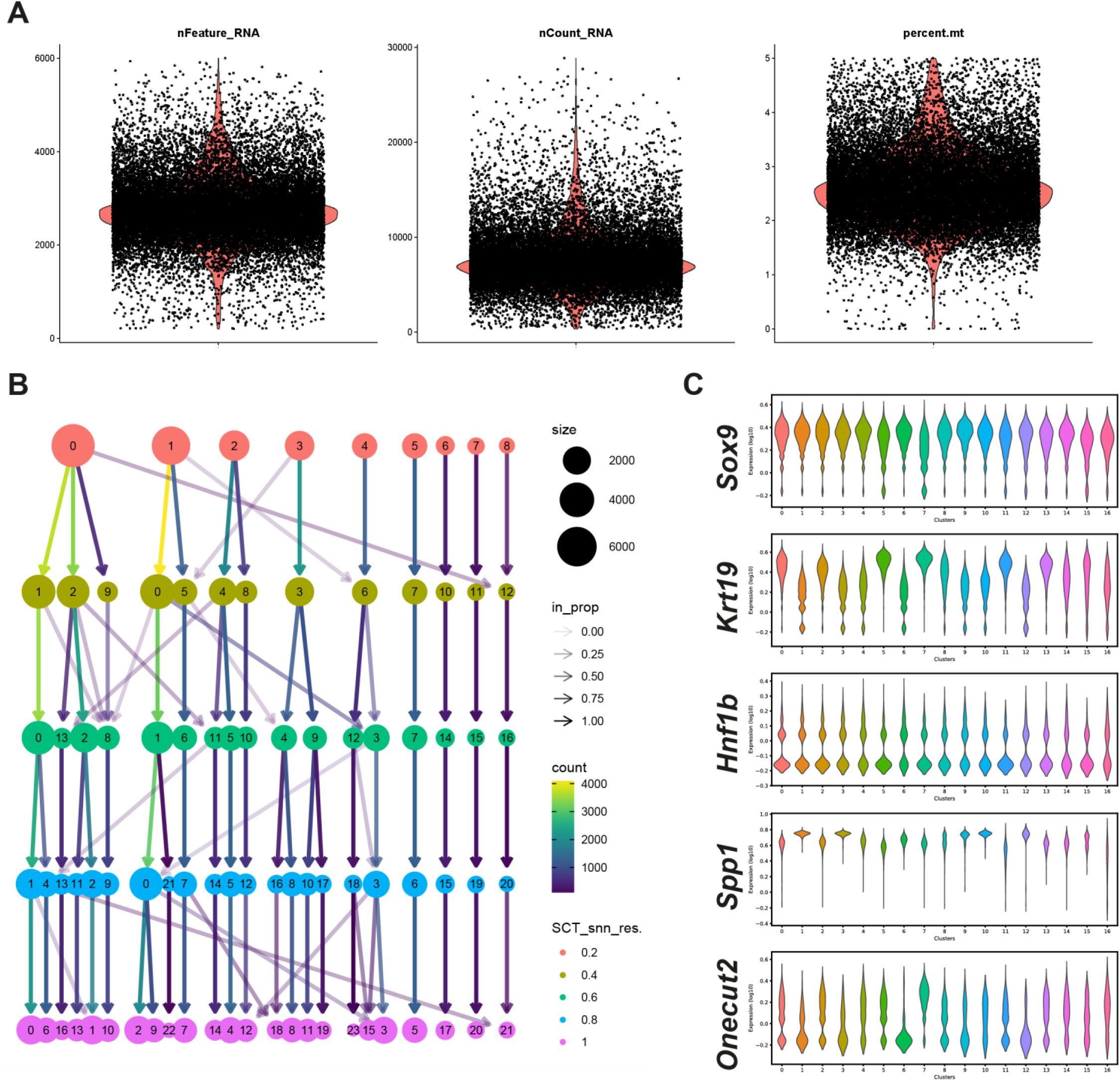

**Supplementary Figure 5.**
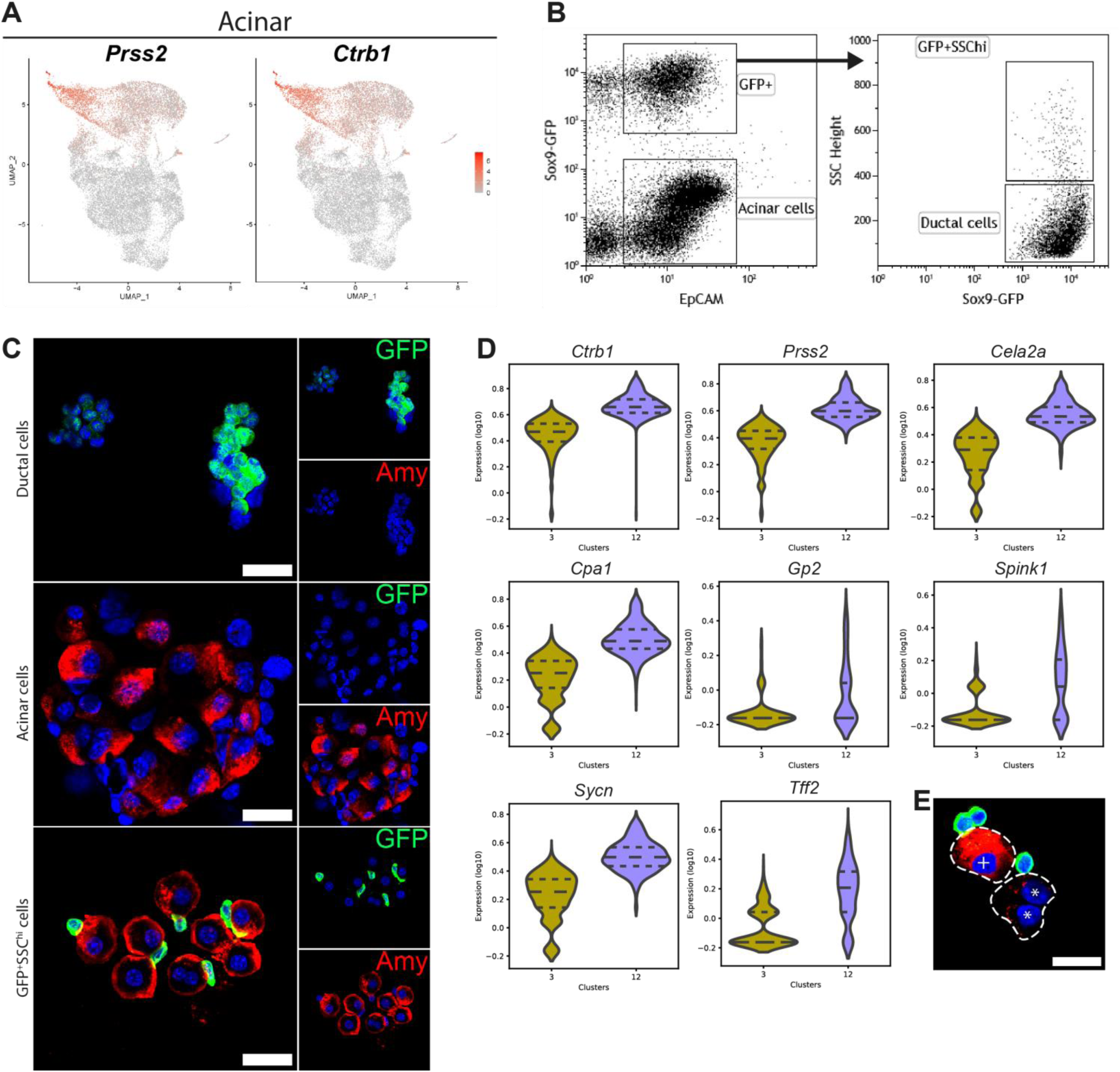

**Supplementary Figure 6.**
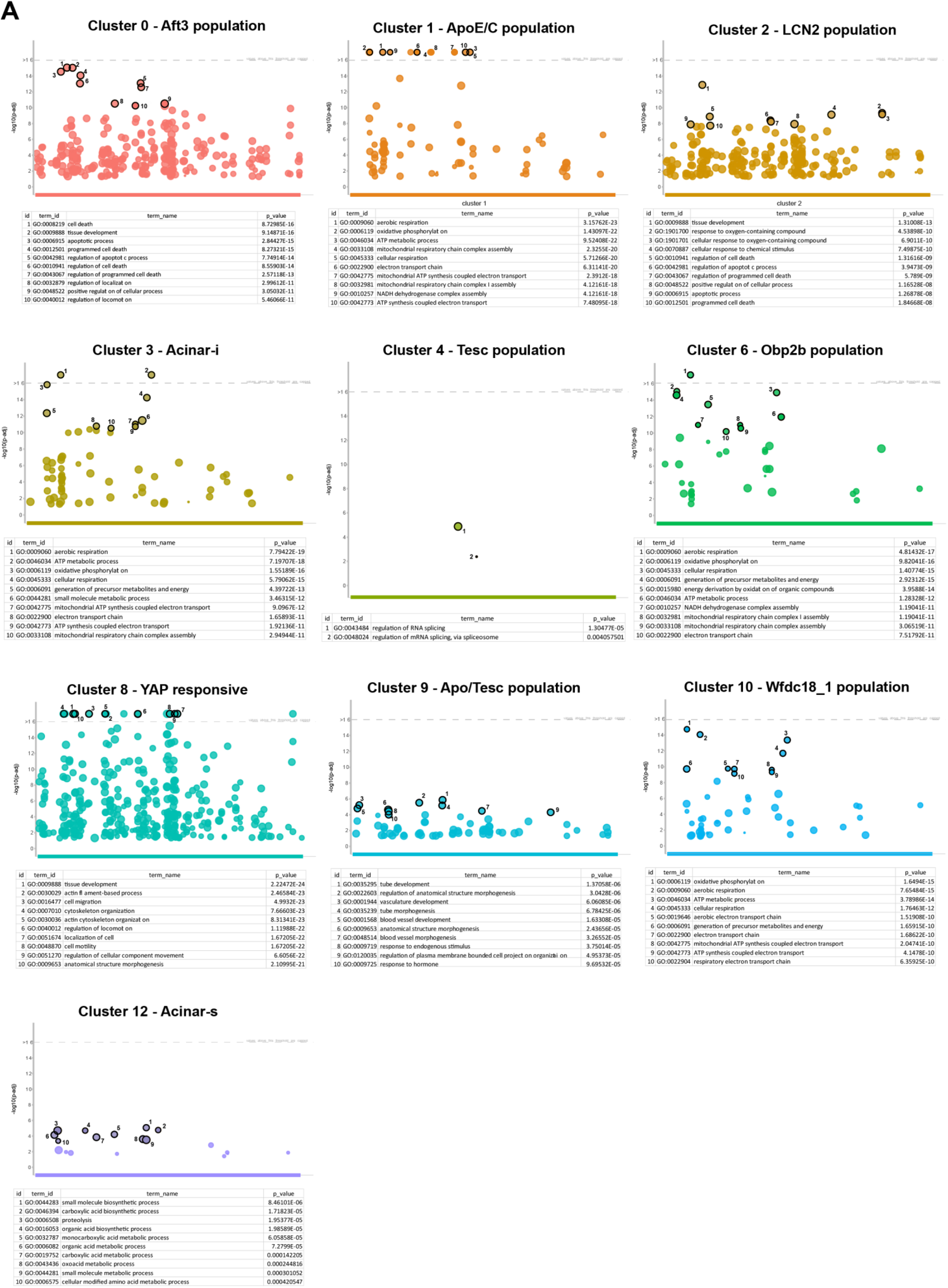

**Supplementary Figure 7.**
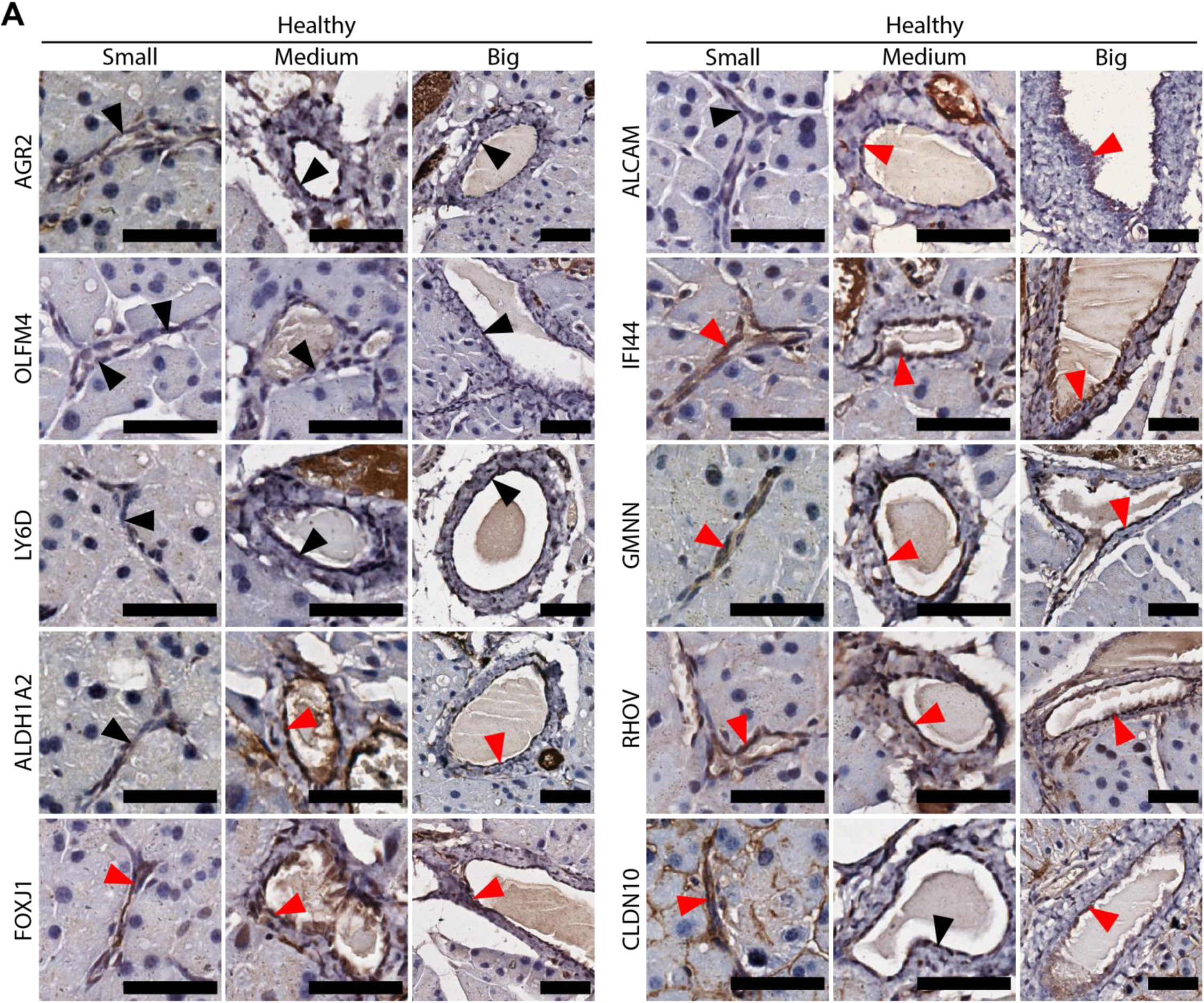

**Supplementary Figure 8.**
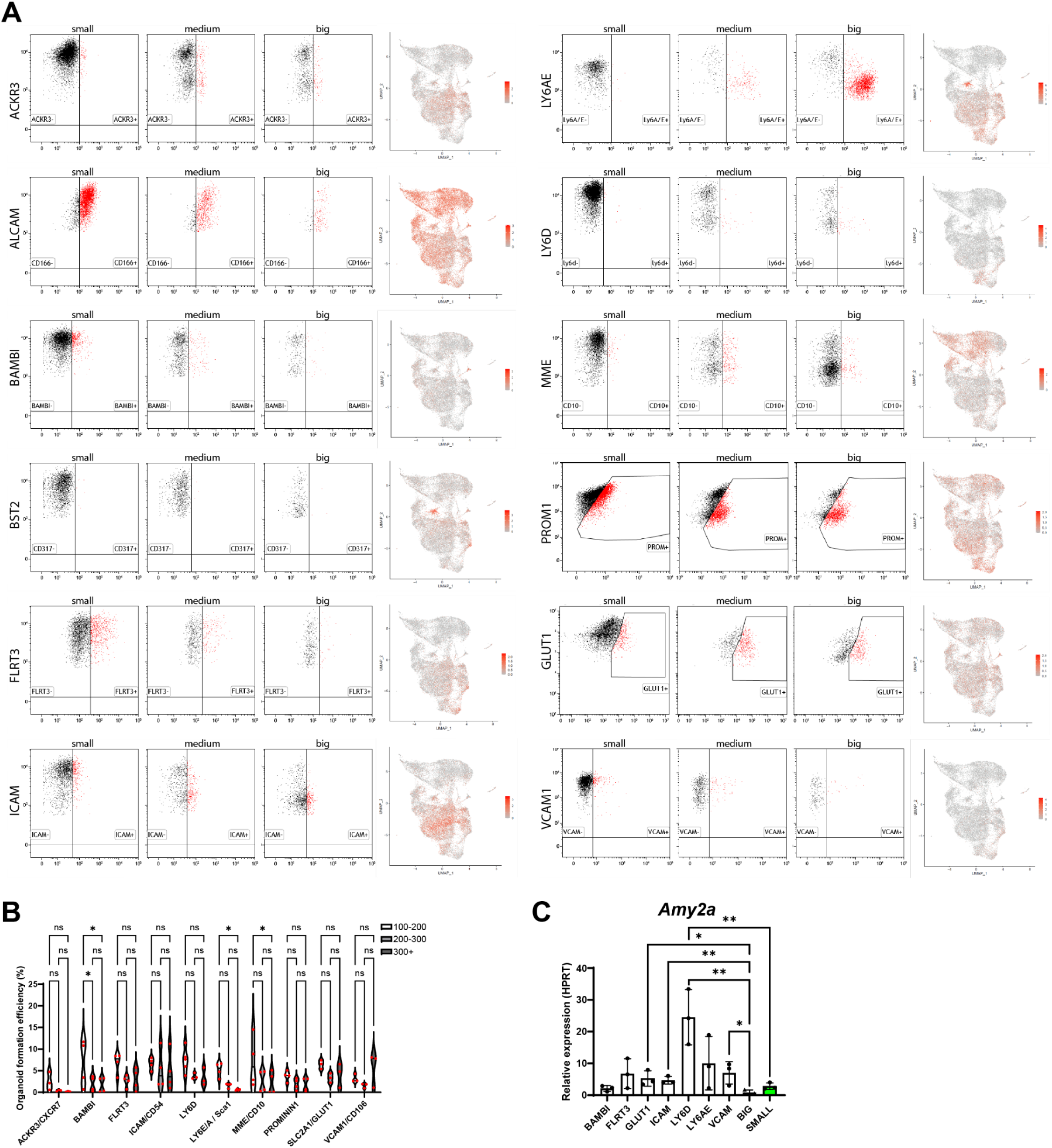

